## Supplementary Material for "LaGrACE: Estimating gene program dysregulation using latent gene regulatory network for biomedical discovery"

**Supplementary Table S1. Clinical characteristics of breast cancer patients vary across clusters.** P-values were calculated with a Kruskal-Wallis test or Wilcoxon rank-sum test for continuous and ordinal variables, or a Chi-squared test for discrete and binary variables.

| | Clinical Feature | All Clusters | Cluster 2 vs 3 | Cluster 4 vs 5 |
|---|---|---|---|---|
| **METABRIC** | **Nottingham histologic index** | **<2.2e-16** | **0.0031** | **0.048** |
| | **Tumor Cellularity** | **2.64E-12** | **0.01732** | **0.01605** |
| | **Inferred Menopausal State** | **6.86E-15** | **0.0002635** | **0.00964** |
| **SCAN-B** | **Nottingham histologic grade** | **<2.2e-16** | **<2.2e-16** | |
| | **KI67 status** | **<2.2e-16** | **<2.2e-16** | |

**Supplementary Table S2. Nottingham histologic grade (NHG) was stratified by clusters on SCAN-B datasets.** G2 is equivalent to 6~7 Nottingham histologic index (NPI), and G3 is equivalent to 8~9 NPI.

| NHG | G2 | G3 |
|---|---|---|
| 0 | 336 | 369 |
| 1 | 768 | 2 |
| 2 | 498 | 9 |
| 3 | 122 | 75 |
| 4 | 2 | 8 |

**Supplementary Table S3. Discrimination indexes of cox proportional hazard model on survival and recurrent events**

| Input | Task | C-index | CPE |
|---|---|---|---|
| Molecular Subtype | METABRIC Survival | 0.628 (se = 0.013) | 0.606 (se = 0.011) |
| **LaGrACE** | | **0.642 (se = 0.013)** | **0.630 (se = 0.012)** |
| ssGSEA | | 0.629 (se = 0.013) | 0.612 (se = 0.012) |
| Pathifier | | 0.606 (se = 0.014) | 0.587 (se = 0.013) |
| Molecular Subtype | METABRIC Distant Relapse | 0.618 (se = 0.014) | 0.600 (se = 0.012) |
| **LaGrACE** | | **0.654 (se = 0.013)** | **0.634 (se = 0.012)** |
| ssGSEA | | 0.630 (se = 0.013) | 0.612 (se = 0.013) |
| Pathifier | | 0.604 (se = 0.014) | 0.585 (se = 0.013) |
| Molecular Subtype | METABRIC Local Relapse | 0.614 (se = 0.021) | 0.583 (se = 0.018) |
| **LaGrACE** | | **0.626 (se = 0.019)** | **0.597 (se = 0.020)** |
| ssGSEA | | 0.622 (se = 0.021) | 0.589 (se = 0.020) |
| Pathifier | | 0.616 (se = 0.021) | 0.592 (se = 0.020) |
| Molecular Subtype | SCANB Survival | 0.561 (se = 0.020) | 0.557 (se = 0.016) |
| **LaGrACE** | | **0.617 (se = 0.019)** | **0.622 (se = 0.018)** |

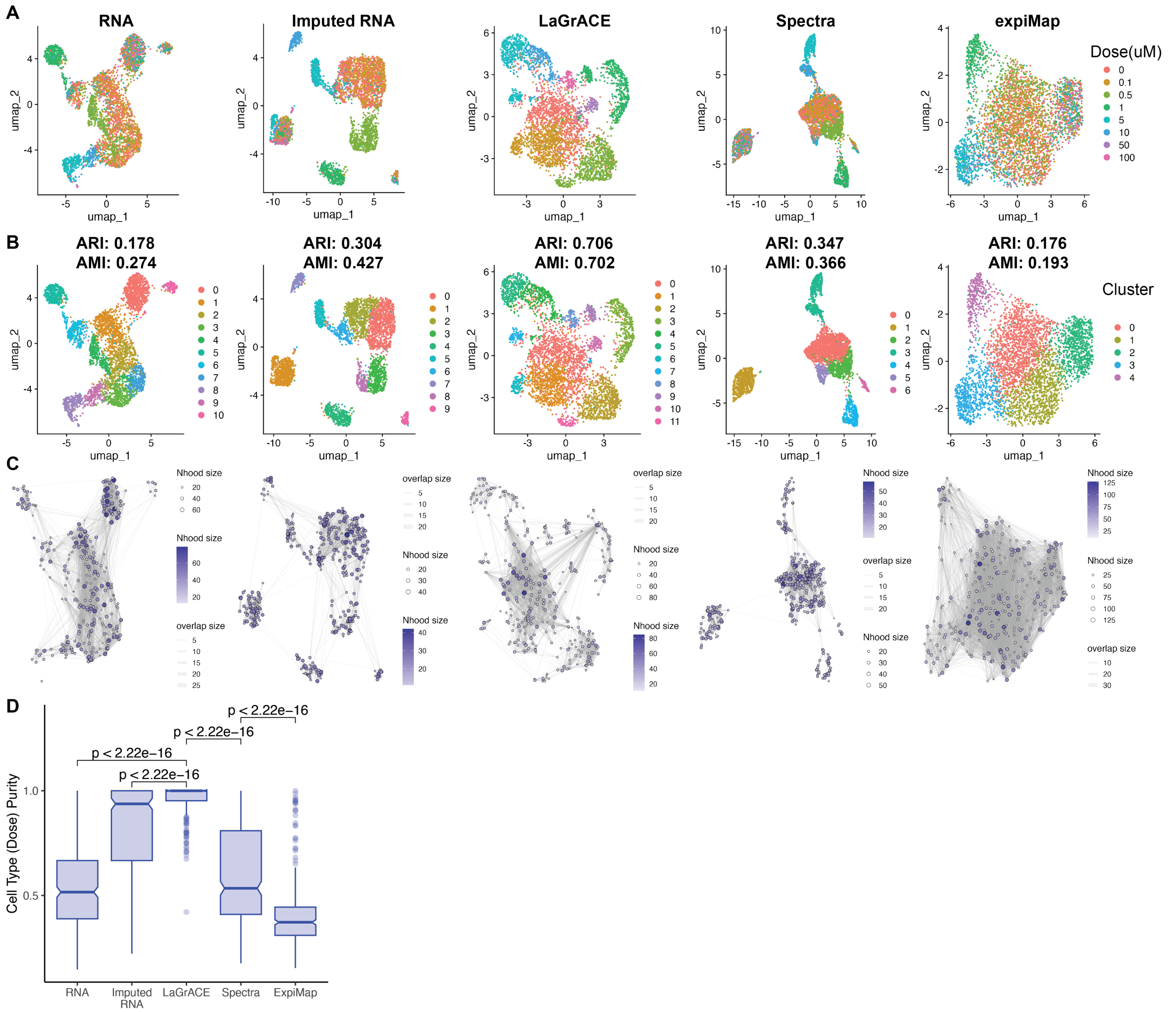


**Supplementary Figure S1. LaGrACE captures dose-response signal from BMS-345541 treatment at single-cell resolution.** A549 lung adenocarcinoma cells were treated with BMS345541 (an inhibitor of nuclear factor κB–dependent transcription) for 24 hours.

(A) A549 cells visualized and colored by dose on UMAP embeddings computed based on unimputed RNA profiles, RNA profiles imputed by SCVI, LaGrACE features and gene set scores inferred by Spectra and ExpiMAP.

(B) UMAP plot of A549 cells colored by clusters based on unimputed RNA (ARI: 0.178, AMI: 0. 274), imputed RNA (ARI: 0.304, AMI: 0.427), LaGrACE features (ARI: 0.706, AMI: 0.702), Spectra gene set scores (ARI: 0.347, AMI: 0.366), and ExpiMAP gene set scores (ARI: 0.176, AMI: 0.193).

(C) Neighborhood graph constructed based on unimputed RNA profiles, RNA profiles imputed by SCVI, LaGrACE features and gene set scores inferred by Spectra and ExpiMAP using Milo.

(D) Boxplot of cell type purity score for single cell Neighborhoods constructed using Milo.

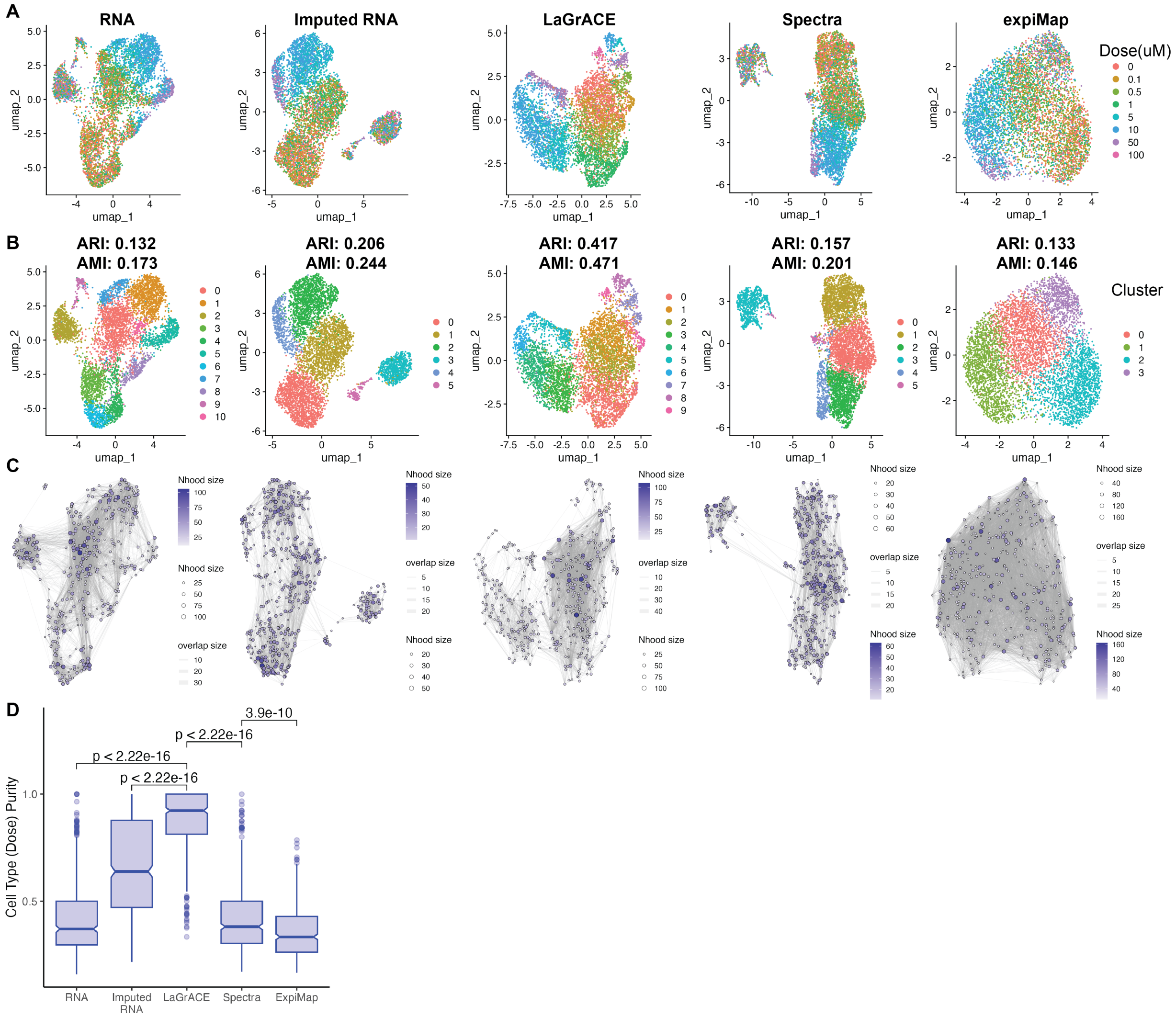


**Supplementary Figure S2. LaGrACE captures dose-response signal from nutlin-3a treatment at single-cell resolution.** A549 lung adenocarcinoma cells were treated with nutlin-3a (a p53-Mdm2 antagonist) for 24 hours.

(A) A549 cells visualized and colored by dose on UMAP embeddings computed based on unimputed RNA profiles, RNA profiles imputed by SCVI, LaGrACE features and gene set scores inferred by Spectra and ExpiMAP.

(B) UMAP plot of A549 cells colored by clusters based on unimputed RNA (ARI: 0.132, AMI: 0.173), imputed RNA (ARI: 0.206, AMI: 0.244), LaGrACE features (ARI: 0.417, AMI: 0.471), Spectra gene set scores (ARI: 0.157, AMI: 0.201), and ExpiMAP gene set scores (ARI: 0.133, AMI: 0.146).

(C) Neighborhood graph constructed based on unimputed RNA profiles, RNA profiles imputed by SCVI, LaGrACE features and gene set scores inferred by Spectra and ExpiMAP using Milo.

(D) Boxplot of cell type purity score for single cell Neighborhoods constructed using Milo.

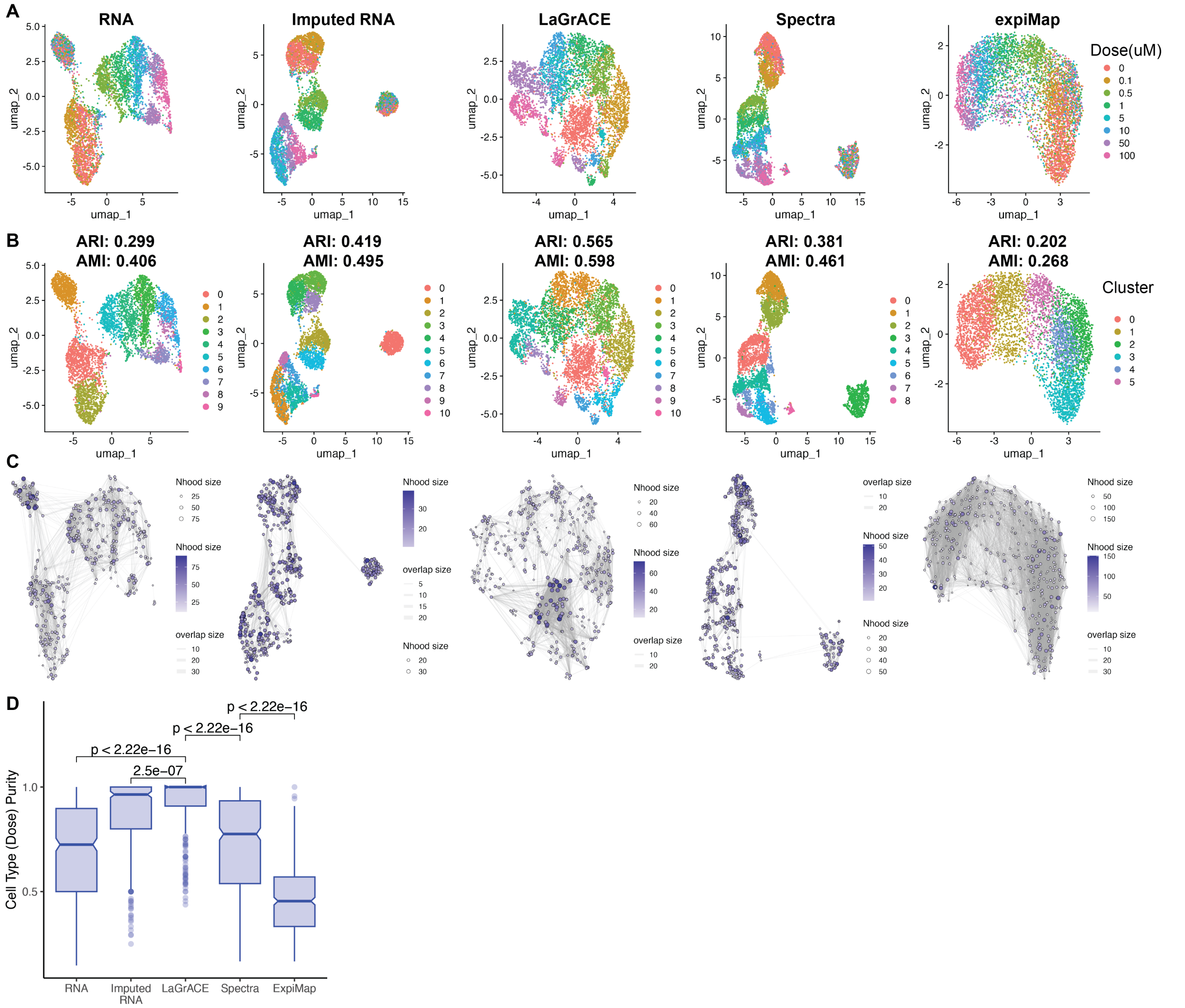


**Supplementary Figure S3. LaGrACE captures dose-response signal from vorinostat treatment at single-cell resolution.** A549 lung adenocarcinoma cells were treated with suberoylanilide hydroxamic acid (SAHA), an HDAC inhibitor, for 24 hours.

(A) A549 cells visualized and colored by dose on UMAP embeddings computed based on unimputed RNA profiles, RNA profiles imputed by SCVI, LaGrACE features and gene set scores inferred by Spectra and ExpiMAP.

(B) UMAP plot of A549 cells colored by clusters based on unimputed RNA (ARI: 0.299, AMI: 0.406), imputed RNA (ARI: 0.419, AMI: 0.495), LaGrACE features (ARI: 0.565, AMI: 0.598) , Spectra gene set scores (ARI: 0.381, AMI: 0.461), and ExpiMAP gene set scores (ARI: 0.202, AMI: 0.268).

(C) Neighborhood graph constructed based on unimputed RNA profiles, RNA profiles imputed by SCVI, LaGrACE features and gene set scores inferred by Spectra and ExpiMAP using Milo.

(D) Boxplot of cell type purity score for single cell Neighborhoods constructed using Milo.

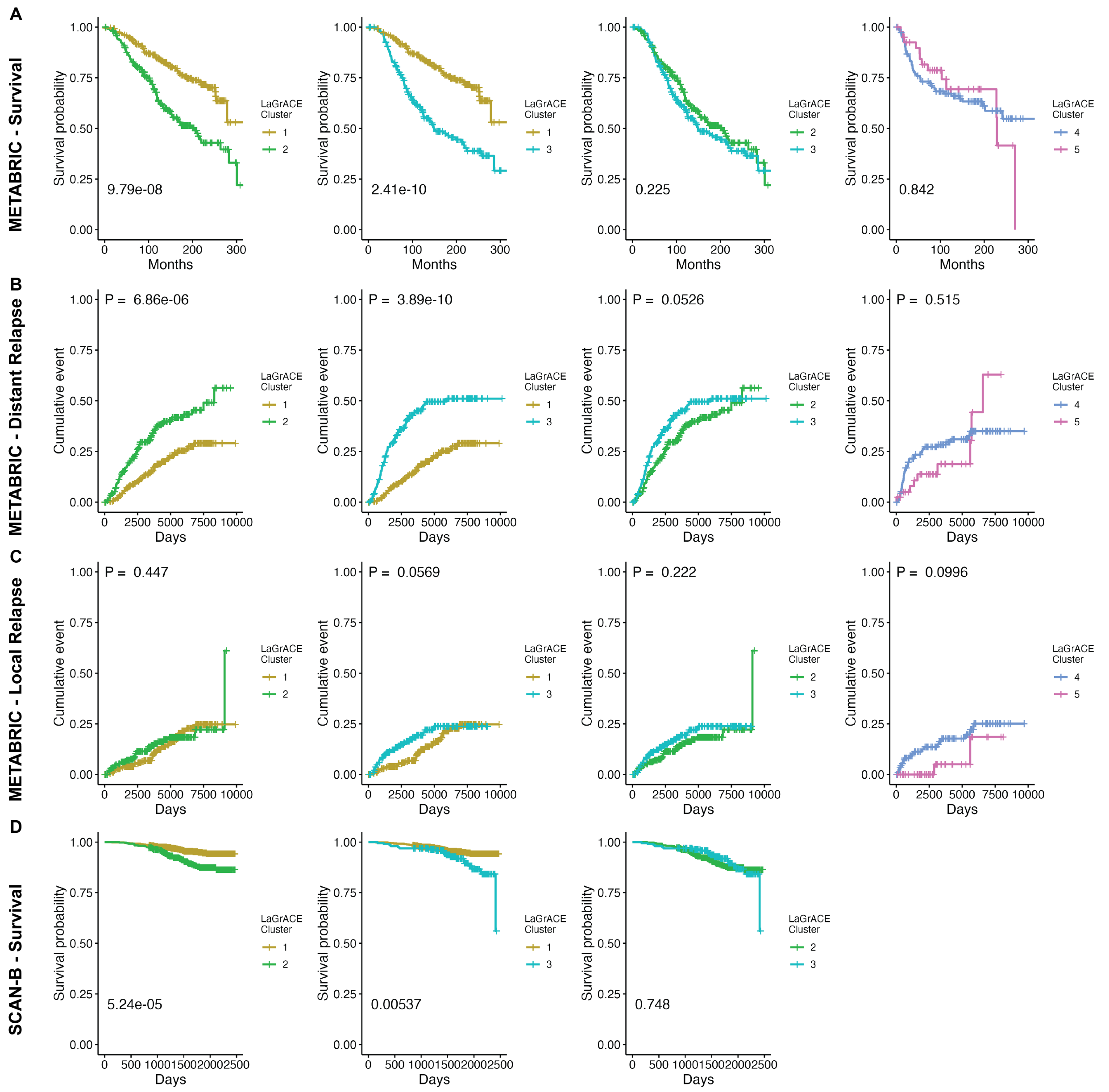


**Supplementary Figure S4. Breast Cancer subject survival analysis and recurrent event analysis for clusters with similar molecular subtype composition on two breast cancer datasets.** Luminal samples were grouped into cluster 1, 2 & 3; Claudin-low samples were grouped into clusters 4 & 5.

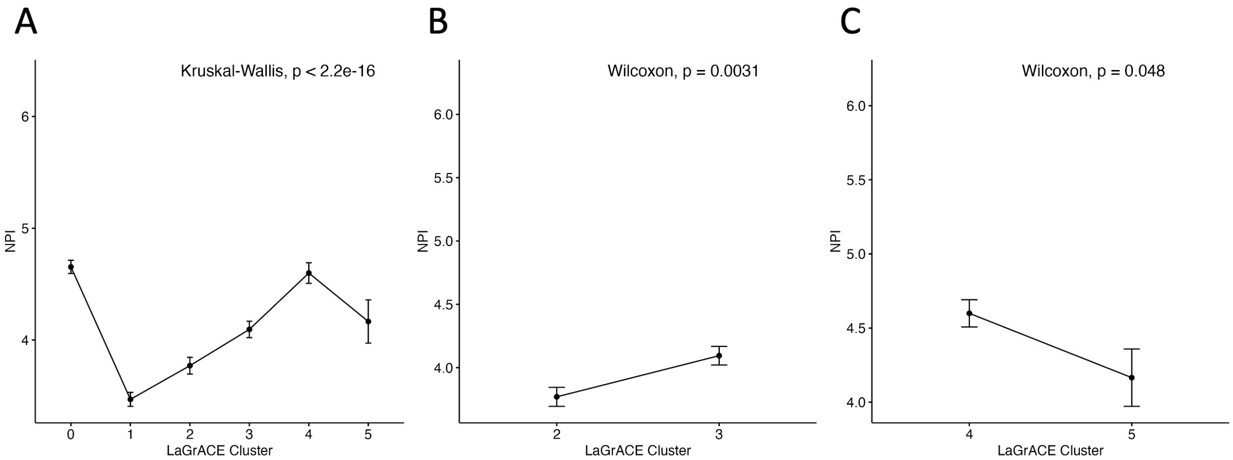


**Supplementary Figure S5. Line plot of Nottingham Prognostic index (NPI) among clusters identified on METABRIC dataset.** Kruskal-Wallis test or Wilcoxon rank-sum test was performed for (A) all clusters, (B) cluster 2 and cluster 3, (C) Cluster 4 and cluster 5

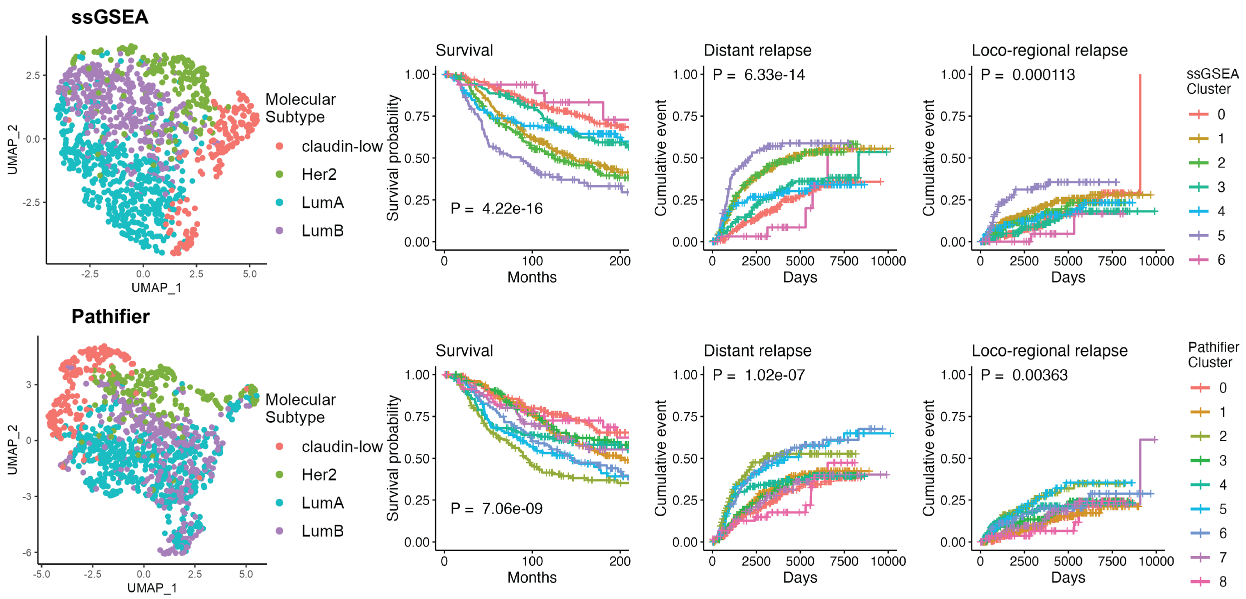


**Supplementary Figure S6. Separation of METABRIC breast cancer samples and Kaplan-Meier estimate of survival and recurrent event by ssGSEA's and Pathifier's clusters**

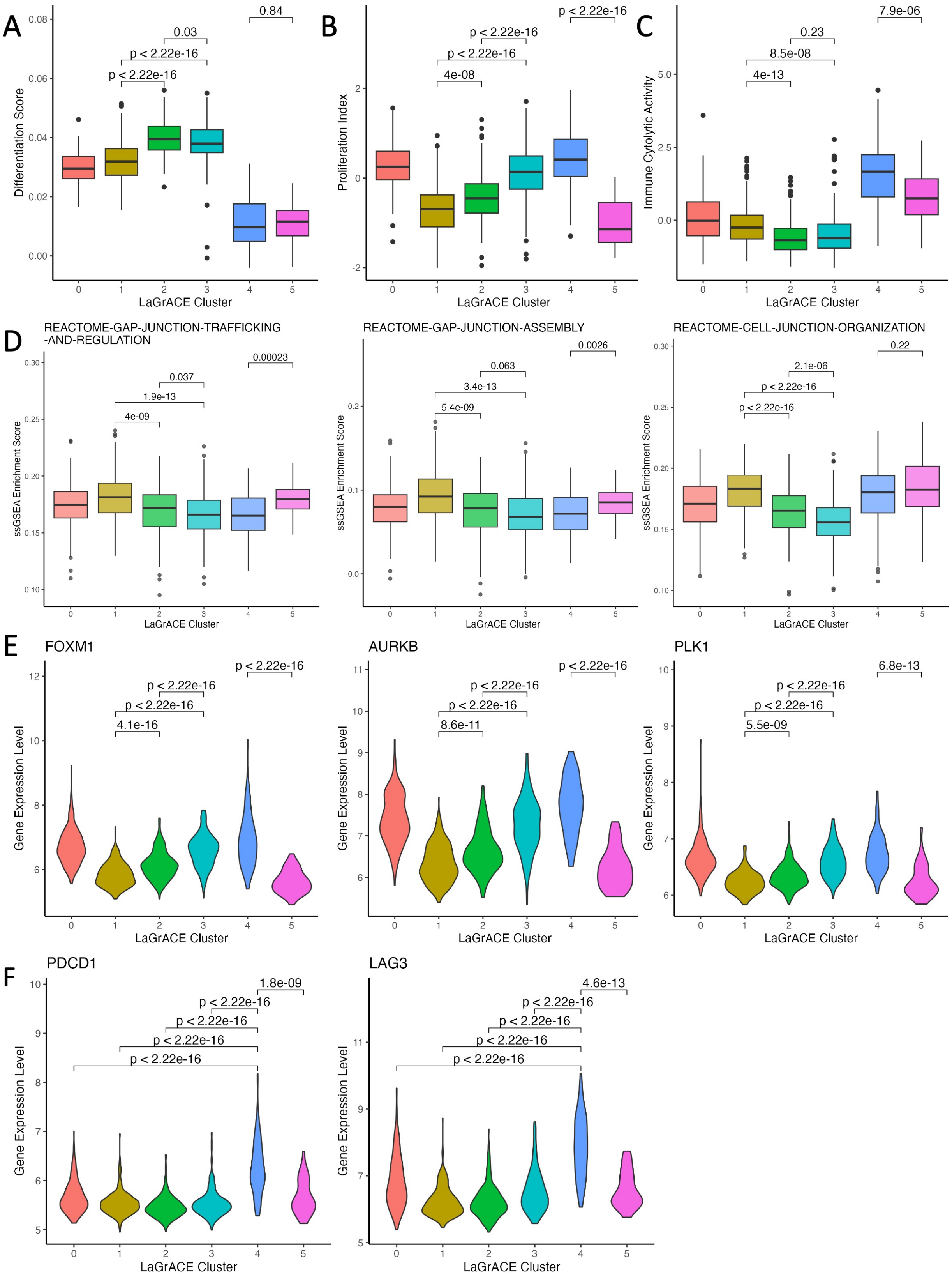


**Supplementary Figure S7. Biological process and molecular mechanism difference among identified clusters in breast cancer.** Box plots of (A) differentiation, (B) proliferation and (c) immune cytolytic scores of each sample,

(D) ssGSEA pathway enrichment score of junction-associated pathways; Violin plots visualizing gene expression of (E) FOXM1, AURKB and PLK1 and (F) immune exhaustion markers, PDCD1 and LAG3. Wilcoxon rank-sum test was conducted to calculate P values.

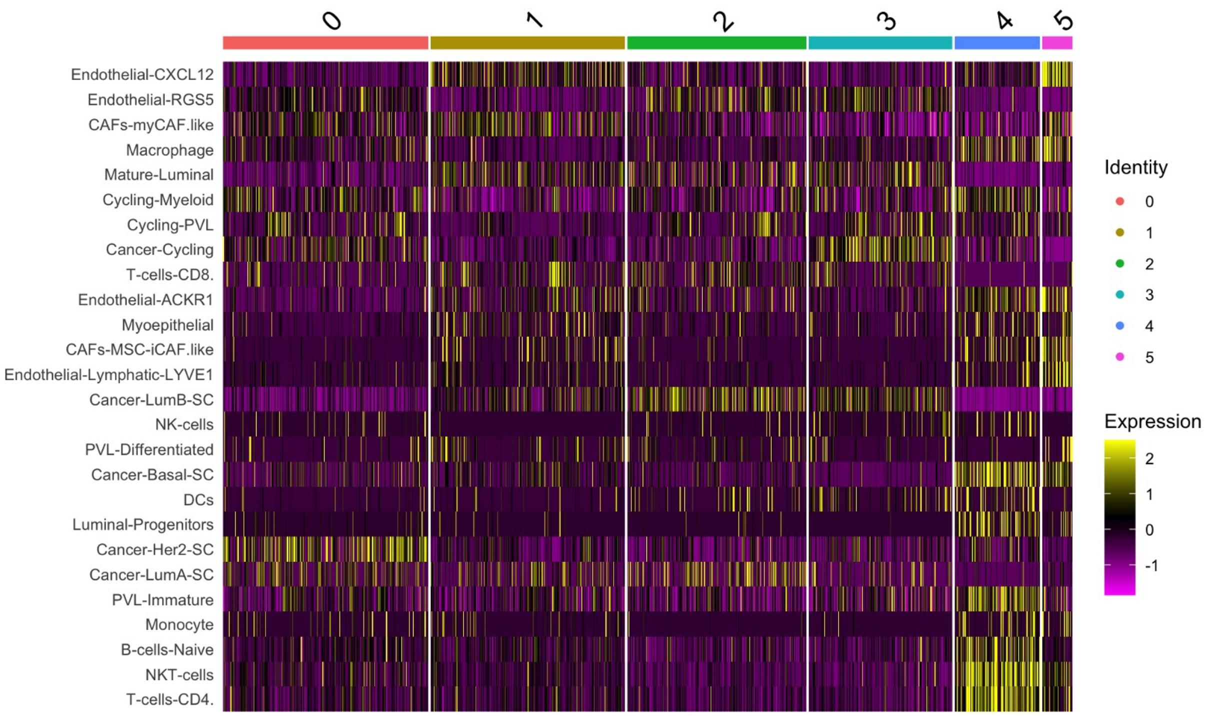


**Supplementary Figure S8. Heatmap of scaled predicted cell type fractions inferred using CIBERSORTx.**

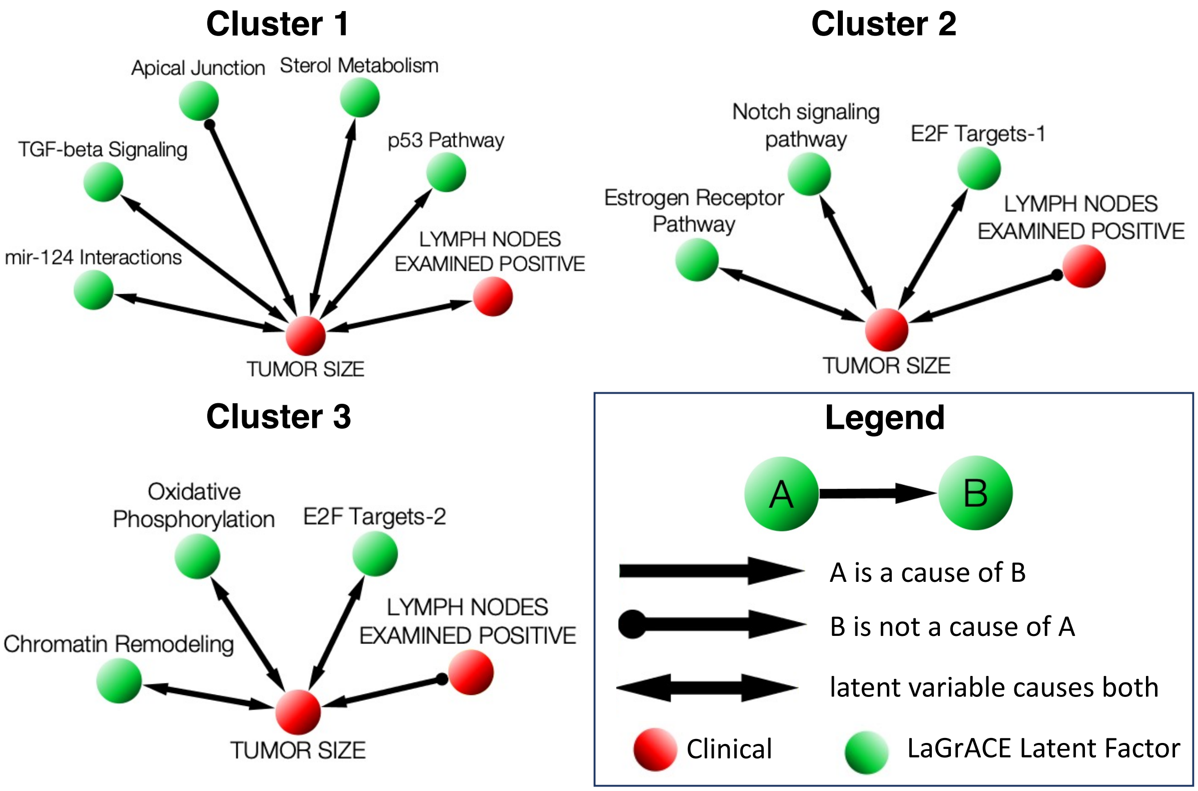


**Supplementary Figure S9. The cluster specific causal graphs for luminal clusters (1, 2 & 3).** Note that besides the edges represented by a direct arrow (A→B), all other edges do not exclude the possibility of a latent confounder. AGE AT DIAGNOSIS: Age of patient at cancer diagnosis (years); LYMPH NODEs Examined Positive: Number of lymph nodes where tumor cells are seen: 0, 1 to 3, 4 to X; TUMOR SIZE: Log2 size of tumor (cm)

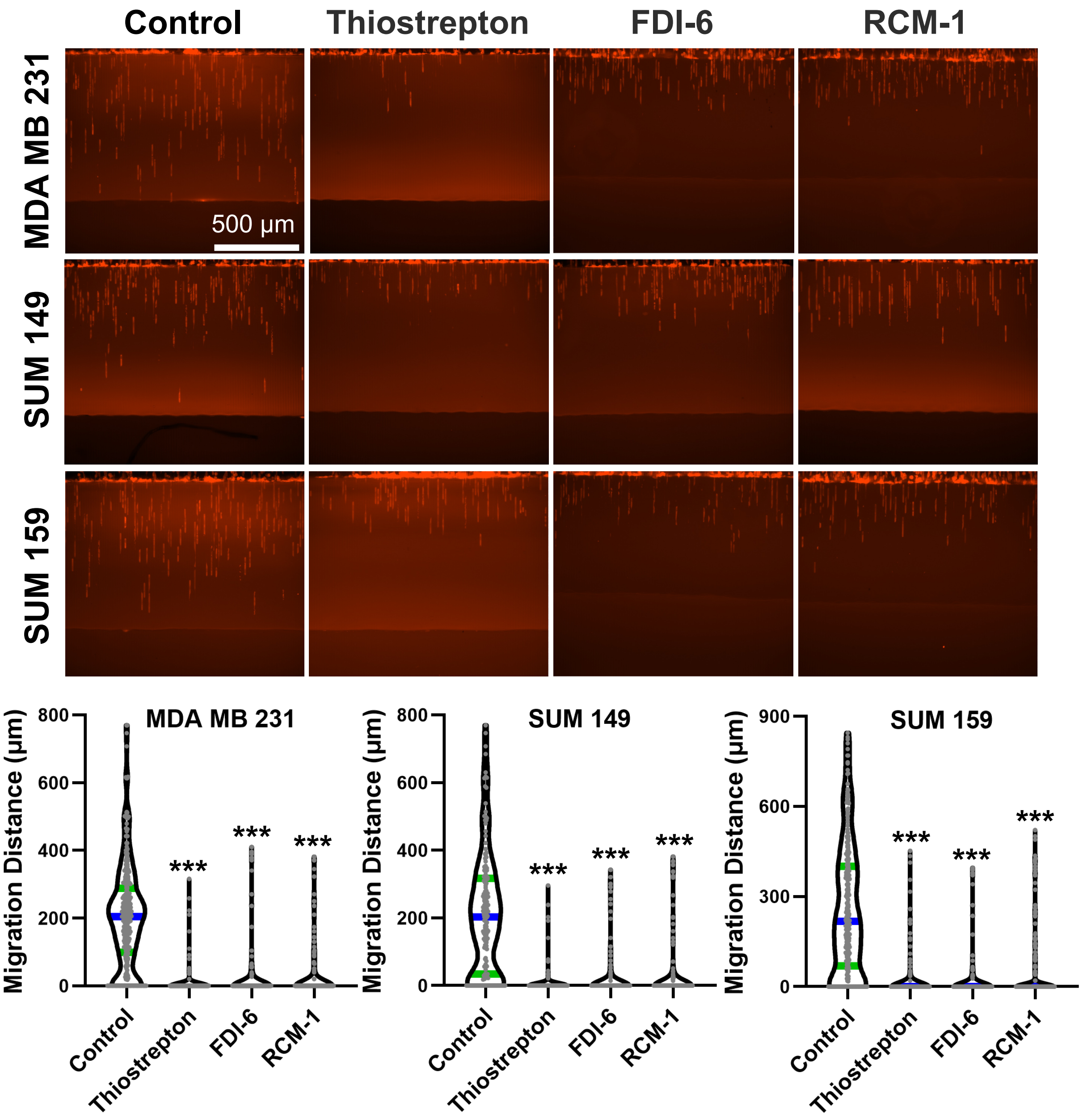


**Supplementary Figure S10. FOXM1 inhibitors inhibit the cell migration of breast cancer cells. (Scale bar: 500 μm)** Each dot represents the cell migration distance in a channel. The green bar represents the median, and the blue bars represent the quartiles. (n = 200 channels). *** refers to P < 0.001 (compared with control).

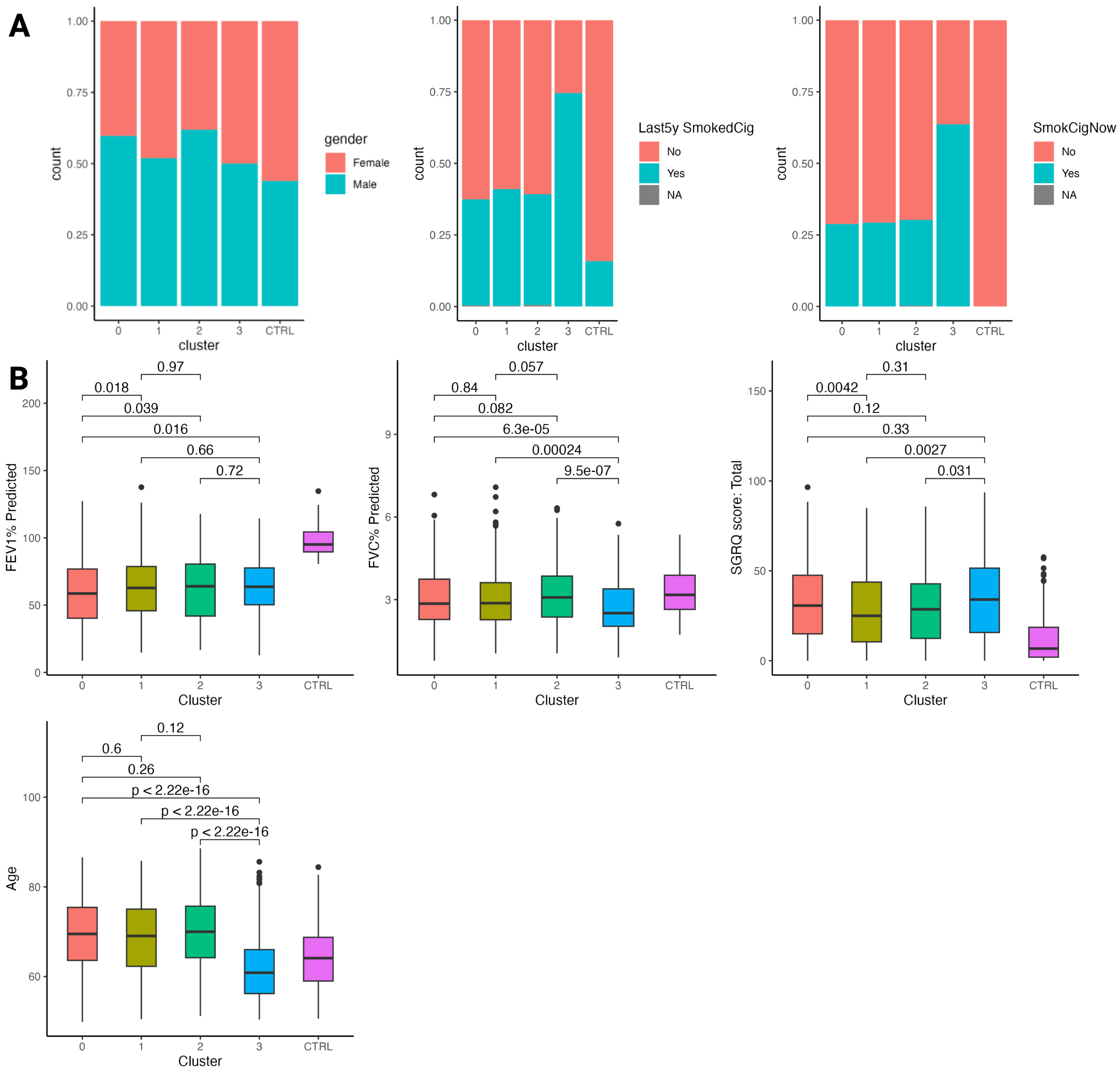


**Supplementary Figure S11. Association between LaGrACE cluster and clinical variables.** (A) Box plots illustrating categorical variables: Gender, Current Smoking Status, and Smoking Status over the Past 5 Years. (B) Box plots depicting continuous variables including Predicted FEV1, Predicted FVC, SGRQ Score, and age. Wilcoxon rank-sum test was conducted to calculate P values.

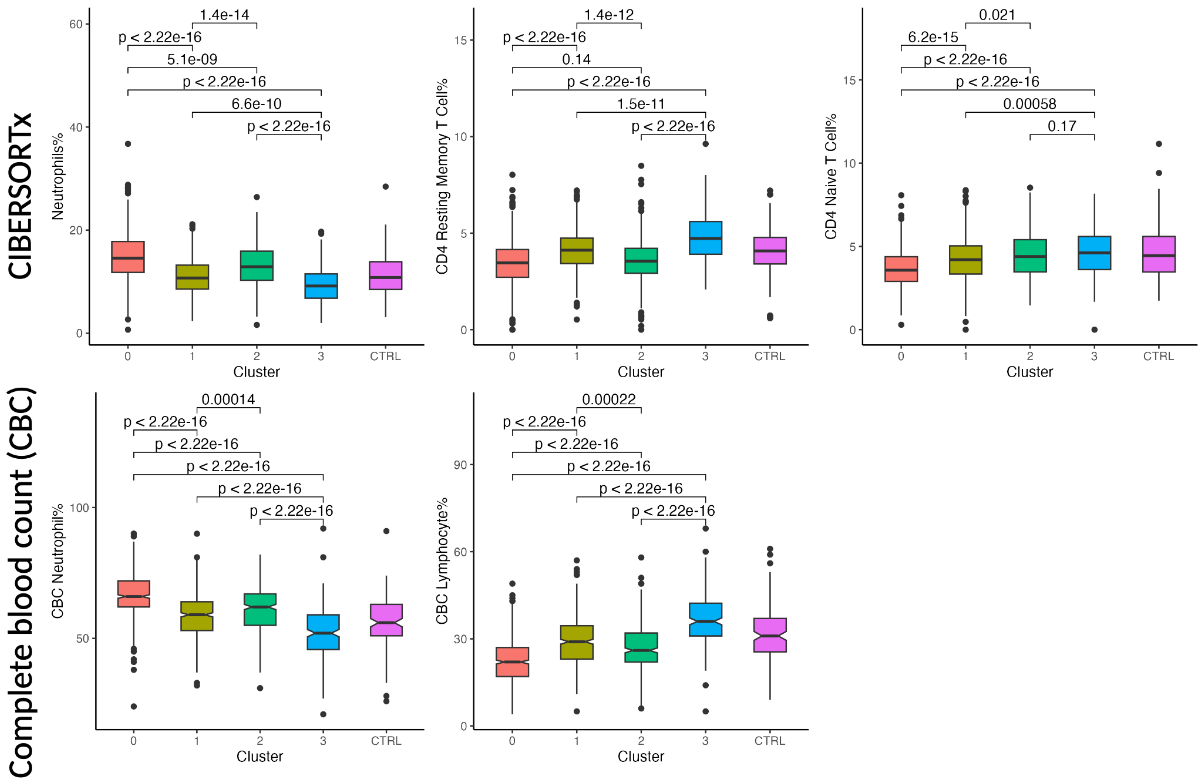


**Supplementary Figure S12. Association between LaGrACE cluster and white blood cell type fraction.** Wilcoxon rank-sum test was conducted to calculate P values.

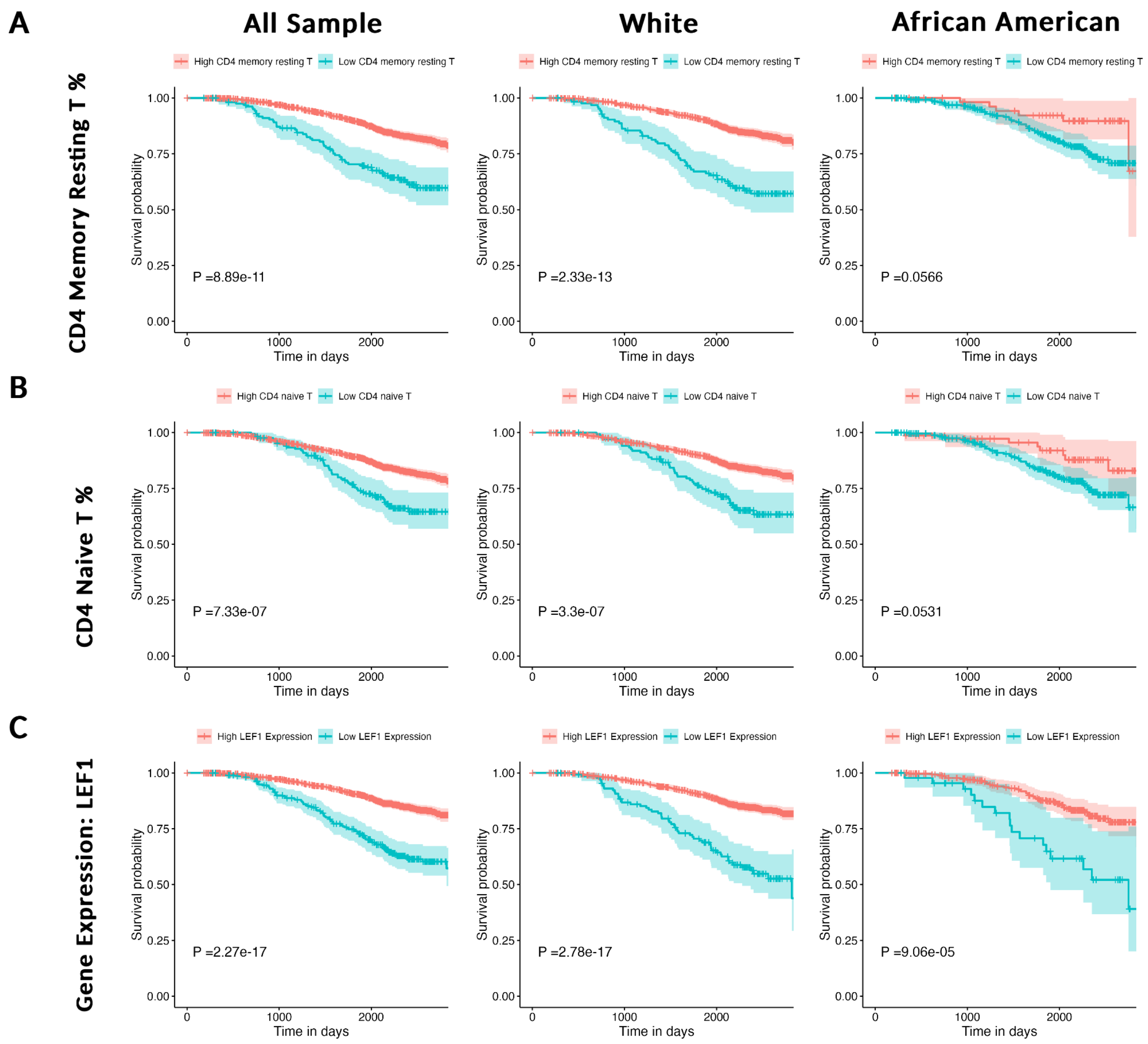


**Supplementary Figure S13: Kaplan-Meier survival curve analysis for COPD patients.** This figure presents the estimation of survival curves based on three different criteria: (A) the fraction of CD4+ memory resting T cells, (B) the fraction of CD4+ naïve T cells, and (C) the expression levels of the LEF1 gene. An optimal cutpoint was determined for each combination of feature and race.

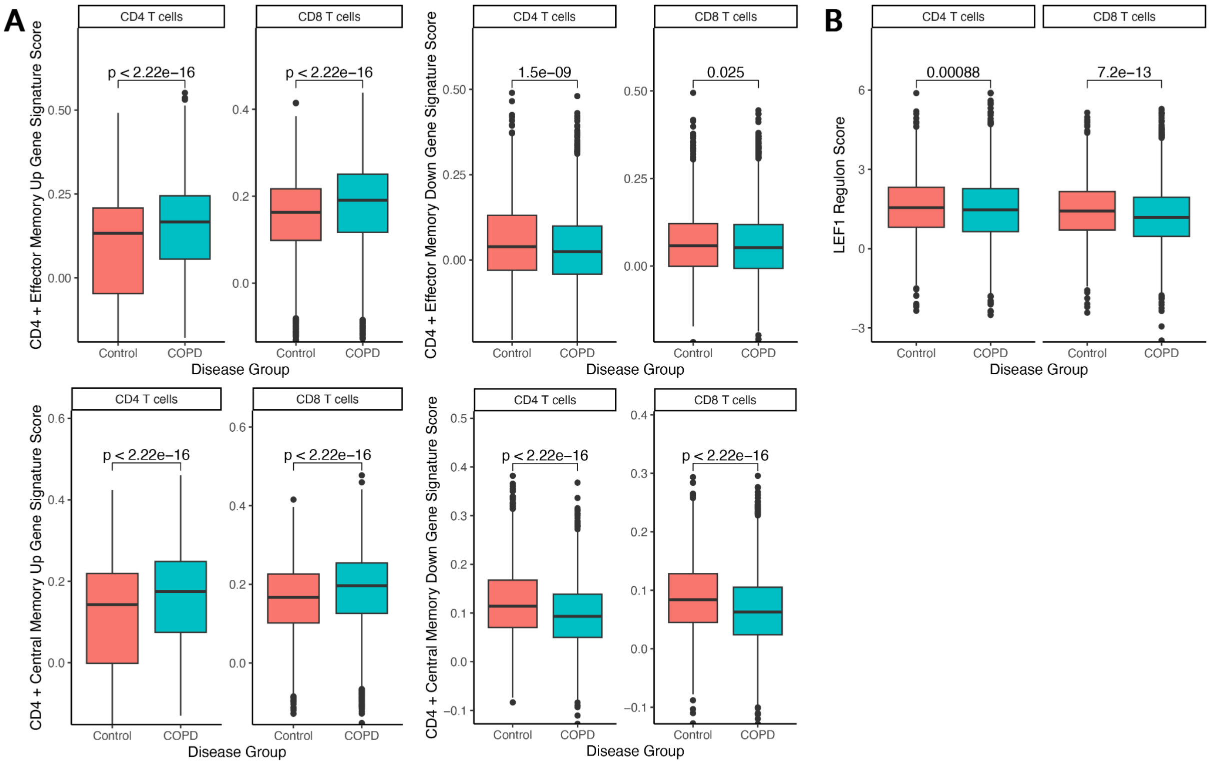


**Supplementary Figure S14: COPD module score and LEF1 regulon score in CD4 and CD8 T cells from human lung tissue.** (A) The COPD module score, calculated based on COPD associated differentially expressed genes identified in memory CD4 T cells from blood. (B) The LEF1 regulon score. For an accurate assessment, the analysis is restricted to samples from individuals aged 55 years and older (10 control and 17 COPD samples).
